# Promising protocol for *in vivo* experiments with betulin

**DOI:** 10.1101/2025.05.12.653392

**Authors:** Pavel Šiman, Aleš Bezrouk, Alena Tichá, Hana Kozáková, Tomáš Hudcovic, Otto Kučera, Mohamed Niang

## Abstract

**Background/Objectives:** Betulin is a promising agent in many areas of medicine and is being investigated, particularly in the field of cancer. However, in *in vivo* experiments, its water insolubility becomes a significant obstacle. This study describes a promising method for the administration of betulin in *in vivo* experiments and the determination of betulin levels in organ samples.

**Methods:** Betulin is dissolved first in ethanol, and this solution is introduced into acylglycerols, followed by evaporation of the ethanol. Olive oil and food-grade lard were determined to be suitable lipids for noninvasive application *per os*. A method for processing the organs of experimental animals for betulin determination was developed. Determination in blood is also likely the only viable option to be used in future clinical studies and practice.

**Results:** The maximum amount of betulin usable (i.e., absorbable by organisms) in olive oil (10 mg/ml), suppository mass (6 mg/ml), food lard (4 mg/ml), and cocoa butter (2 mg/ml) carriers was found microscopically. A specific distribution of betulin concentration in the organs of experimental animals (Wistar rats) after a weekly diet containing betulin was discovered. The blood was shown to be particularly advantageous, as it allows continuous monitoring of betulin levels in the body. In these pilot experiments, a statistically significant (P < 0.001) synergistic effect of betulin on solid Ehrlich adenocarcinoma tumors was observed when betulin was combined with cytostatic Namitecan (NMRI mice). The high-purity betulin used in this study is very stable even under fluctuating storage conditions.

**Conclusions:** Our study proves that both the method of betulin administration and the proposed analytical procedure could greatly increase the reliability and reproducibility of *in vivo* studies and future preclinical and clinical studies on the effects of betulin and possibly other similar water-insoluble triterpenoids on living organisms.

## 1. Introduction

Betulin, a triterpenoid found in considerable amounts in birch bark (Betulaceae), is a promising agent in many areas of medicine and is being investigated, particularly in the field of cancer [1–4]. Betulin has been extensively studied in *in vitro* experiments, where its near insolubility in water is not a major drawback because of the possibility of using excipients, usually dimethyl sulfoxide (DMSO) [5,6]. However, in *in vivo* experiments, its insolubility becomes a significant obstacle, as it cannot be administered with confidence that it is sufficiently available in the body in molecular form to reach therapeutically effective levels. Many attempts have been made to deliver betulin into the bodies of experimental animals, including chemical modifications of betulin, but their results are controversial; e.g., Kuznetsova et al. [7] presented lethal doses for some animals that were quite unlikely to reach such levels of betulin in animals. Betulin, its numerous chemical derivatives [4], betulinic acid, lupeol, or other triterpenoids, are administered to experimental animals in a variety of ways. This is illustrated quite comprehensively in the work of Saneja et al. [8], who described the examples of the administration of betulinic acid in the form of nanoemulsions, liposomes, polymeric or magnetic nanoparticles or polymer conjugates with carriers such as cyclodextrins, polyethylene glycols, biodegradable polymers or even carbon nanotubes. A detailed review of the administration of betulin and its chemical derivatives [9] reported similar methods, with an emphasis on nanoparticulate drug delivery with organic or inorganic nanocarriers. Most of these methods are relatively difficult, expensive and/or do not provide adequate or controllable delivery of the selected triterpenoid to the tissues.

The routes of administration also vary considerably, especially in terms of their efficacy and invasiveness: oral, intravenous, subcutaneous, intramuscular, intraperitoneal, or inhaled. Many studies have not even described the exact procedure used to prepare the form of betulin; sometimes, the relevant citation is not even given but only mentions that a certain amount of betulin was administered orally [10,11], intraperitoneally [12], etc. In such cases, betulin is usually used in the form of a suspension of varying fineness (such as by Zakrzeska et al. [13], e.g.) and not in the form of a true solution. The bioavailability of this form of betulin is very controversial and usually very low [4,9,14,15], and good reproducibility cannot be assumed. Greater bioavailability, but only in extrinsic use, can be achieved by using betulin solution in Oleogel-S10 (a mixture of oils and glyceryl behenate; [16,17]); however, even here, the direct dissolution of betulin is chosen, which does not ensure the perfect molecularity of the solution.

The aim of this study was to develop the simplest noninvasive way of using betulin in experimental animals with respect to its possible future simple, inexpensive and reproducible use in medicine. Moreover, we designed a simple method of processing animal tissues to determine the concentration of betulin in biological samples, i.e., to determine the effective concentration in the organism.

## 2. Materials and Methods

### 2.1. Materials and equipment

#### 2.1.1. Animals

NMRI mice (35 males used for the final experiments that provided the resulting data + 15 males used for the optimization of individual methodologies (e.g., administration processes, experiments with clotted blood) without relevant experimental data, weighing 32–40 g) and Wistar rats (2 males, weighing 233 and 245 g) were all fed standard laboratory mouse/rat chow (Velaz, Czech Republic) and water ad libitum, under laboratory conditions, on a 12:12-h light-dark cycle.

#### 2.1.2. Drugs

Betulin (99.7%) was prepared according to the improved method of Šiman et al. [5] (see Section 2.5 below). Other materials included lupeol (> 94%, Sigma-Aldrich, St. Luis, MO, USA), Namitecan (Sigma-Aldrich, USA), olive oil (pharmaceutical grade), suppository mass (pharmaceutical grade), food grade lard, food grade cocoa butter, ethanol (96%, VWR, France), NaOH (Sigma-Aldrich), and chloroform (Sigma-Aldrich).

#### 2.1.3. Chromatography

Gas chromatography with mass spectrometry (GC-MS) detection was chosen for quantification of the isolated triterpenes (Šiman et al. [5], Caligiani et al. [18], Hassan et al. [19]). The use of the GC-MS method was given by available equipment and established practices, but liquid chromatography methods can be significantly more sensitive (e.g., UHPLC-MS; Rathod et al. [20]). The estimated sensitivity of the GC-MS used in our experiment was significantly below 1 μg/g of sample.

The samples were derivatized with a mixture of pyridine (Sigma Aldrich, USA) with N,O-bis(trimethylsilyl)trifluoroacetamide (Supelco, USA) (1:1 v/v) at 45 min at 75 °C. The trimethylsilyl derivatives of the triterpenes were determined via gas chromatography-mass spectrometry (Agilent Technologies—GC 7890A, MS 7890A, USA) and capillary column J and WDB-5 MS 60 m × 250 μm × 0.25 μm (Agilent Technologies, USA). The injector temperature was 280 °C, and the oven was programmed as follows: initial temperature, 70 °C; hold time, 1 min; and rate, 15 °C/min to 300 °C. The pressure at the column head was 50 kPa. The mass spectrometer was used in electron impact mode (electron energy 70 eV, source temperature 230 °C and quadrupole 150 °C). Sample concentrations were quantified via the external standard method, which is resistant to derivatization differences. The ratios of betulin to lupeol were quantified via calculation of the calibration curve.

#### 2.1.4. Microscopy

Microscopically, the maximal concentration of betulin in the carriers was determined via a Nikon Eclipse 90i microscope with a Plan Apo 40x DIC M N2 (Nikon Corporation, Tokyo, Japan) and NIS-Elements AR 3.20 software (Laboratory Imaging, Prague, Czech Republic).

#### 2.1.5. Brief description of the betulin preparation

1. Soxhlet extractor was used for extraction from dried outer birch bark in 96% ethanol, and the ethanol extract was dried;
2. The dried extract was dissolved in hot acetone (>40 °C), triterpenoids were precipitated with water, and the precipitate was filtered and dried (after this step, all the products were crystalline, and for subsequent handling with precipitates, filtration was quite sufficient);
3. The precipitate was dissolved in hot ethanol (>40 °C) and betulinic acid, other acidic substances, polyphenols and lipids were removed from the hot solution by adding an ethanolic solution of CaCl_2_ and then an equimolar ethanolic solution of NaOH during intensive stirring. Impurities were removed with the precipitate with Ca(OH)_2_ and NaCl via filtration. The filtrate was concentrated by evaporation up to 1/3–1/4 of the original volume and crude betulin and lupeol were freely crystallized (∽4 °C);
4. Dried crystals from Step 3 were dissolved in chloroform (∽one part of the crystals from Step 3 was dissolved in 16 parts of chloroform (w/v)), and betulin was precipitated by the addition of petroleum ether (lupeol is much more soluble in hydrocarbons than betulin), followed by filtration of the precipitate and washing with petroleum ether. This step was repeated twice;
5. Dried precipitate was dissolved in chloroform and filtered through silica gel (1–2 cm high column), and the filtrate was dried (to yield a snow-white amorphous substance);
6. Pure betulin was crystallized from a hot ethanolic solution via cooling.

### 2.1. Methods

#### 2.2.1. Administration form

Betulin was dissolved in hot ethanol (>60 °C) at a concentration of 20 g/l. This solution was then added in appropriate amounts in batches to the selected carriers (olive oil, suppository mass, food-grade lard, and food-grade cocoa butter) at 80 °C, and the mixture was further heated with stirring until the ethanol was completely evaporated. Attempts were also made to add the entire amount of ethanol solution at one time, with virtually the same result. Since the hot ethanol solution evaporates rapidly in air, care must be taken during addition so that the nascent betulin microcrystals on the solution container do not contaminate the carrier. However, slight foaming of the carrier after the addition of the ethanol solution is not harmful.

To determine the concentration of betulin that can be used in the carriers, the resulting concentrations of 2, 4, 6, 10 and 16 mg betulin per g carrier were chosen. The melted samples (60–80 °C) were assessed microscopically via enriched carriers shortly after preparation and then after storage at 4 °C for three weeks. All images were taken under the same illumination conditions. The resulting focused image was created with a set of successive slices with a step size of 1.2 μm on the Z axis.

#### 2.2.2. Administration of betulin *per os*

The rats were fed a standard diet supplemented with lard containing betulin. The powdered diet was mixed with lard prepared according to the procedure described above with a betulin concentration of 3 g/kg lard to obtain a final dietary betulin concentration of 160 mg/kg. The pellets were prepared from the mixture and made available to the animals ad libitum for 1 week. The animals were then sacrificed, and samples of heparinized blood, spleen, liver, kidney, heart, and adipose tissue were collected to determine the betulin content via the procedure described below.

Twenty-eight mice used in the Namitecan–Betulin experiment (see Section 3.5.) were fed a standard diet, and betulin was applied by gavage as a solution in olive oil on Days 2, 3, 5, and 7 after tumor transplantation. The enriched olive oil was prepared via the method described above at a concentration of 0.5 mg per 1 ml of oil.

#### 2.2.3. Administration of betulin *per rectum*

After overnight starvation, 7 mice were injected with 200 μl of a liquid suppository (37 °C) containing betulin (6 mg/ml) via a catheter *per rectum*. After 2 and 5 h, the mice were sacrificed, and betulin levels in the blood, liver, spleen, jejunum, and duodenum were determined.

#### 2.2.4. Determination of betulin concentrations in organs

Heparinized blood was separated into plasma and cell sediment by centrifugation, and solid organs were roughly homogenized in 1 ml of saline. The amount of processed solid samples did not exceed 1 g, and 0.5 ml of plasma or complete blood (with 1 ml of saline) was always used. All the samples were further processed in glass test tubes to avoid the adsorption of hydrophobic betulin or lupeol at the surface. To all the samples, 4 μg of lupeol dissolved in ethanol (200 μg/l) was added to all the samples as an external standard. Then, 2 ml of a 20% NaOH solution was added, and the mixture was shaken at a temperature of approximately 60 °C until complete dissolution of the tissue was achieved, as reflected by the total transparency of the solution. After the addition of 2 ml of chloroform, the mixture of the two immiscible liquids was shaken vigorously at normal temperature, and each sample was shaken for a total of approximately 5 minutes. The two liquid phases were then separated by centrifugation (1500 × g/10 min), the lighter aqueous phase was carefully removed, and 1 ml of the pure chloroform phase was collected in vials. The vials were left open overnight at laboratory temperature in a fume hood to allow the chloroform to evaporate completely. The samples were further processed and measured via gas chromatography with mass spectrometry detection. Standardization was performed in the same manner using 0.5 ml of intact blood, to which lupeol (4 μg) and betulin (2, 5, 10, 20, 50, and 100 μg) were added as standards for constructing a calibration curve.

Terminological remark: Lupeol is used as an internal standard for GC-MS but is considered an external standard from a biological point of view.

#### 2.2.5 Synergistic effects of Namitecan and Betulin

Twenty-eight mice were subcutaneously injected with a solid form of Ehrlich’s adenocarcinoma, resulting in a total of 106 cells. The mice were randomly divided into 4 groups of 7 animals each: 1. control without betulin and Namitecan, 2. with betulin only, 3. with Namitecan only, and 4. with both betulin and Namitecan. The cytostatic drug Namitecan was injected on Days 3 and 7 after tumor transplantation (15 μg/g body weight) [21], and betulin was applied *per os* by gavage at a dose of 5 μg betulin per 1 g body weight on Days 2, 3, 5, and 7 after tumor transplantation (as a solution in olive oil at a concentration of 0.5 mg/ml). The groups without betulin (1st and 3rd) received an equivalent amount of olive oil without betulin by gavage at the same time. The mice were then left without further treatment for the survival test. The duration of the experiment was 36 days, which corresponds to the survival time of the longest living mouse.

#### 2.2.6. Statistical analysis

The data were statistically evaluated with NCSS 10 statistical software (2015, NCSS, LLC., Kaysville, UT, USA) and MS Excel 2016 (Microsoft Corp., Redmond WA, USA). The normality of the data distribution was tested by the kurtosis normality test and was not rejected. Therefore, the data are presented as the mean and standard deviation of the sample 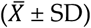. The solid Ehrlich tumor weights after different types of treatment were compared via the 2-sample t test (and Levene’s test was used to check the homoscedasticity assumption). A p value of less than 0.05 was considered statistically significant.

## 3. Results

### 3.1. Administration form

It was proven microscopically that by introducing an ethanol solution of betulin into heated acylglycerols, its true, i.e., molecular, solutions were obtained. Only after a certain concentration was reached could easily recognizable long microcrystals or characteristic ‘hedgehog-like’ crystal clusters with a size of μm begin to fall out. Among the carriers studied, olive oil was the most suitable. It remained as a true solution up to a betulin concentration of 10 mg/ml. Suppository mass could absorb up to 6 mg/ml without precipitating crystals, and food lard could absorb at least 4 mg/ml. Cocoa butter proved unsuitable, as betulin microcrystals appeared even at a concentration of 2 mg/ml. Storage at 4 °C for one month revealed that the true solutions were relatively stable; after repeated melting by heating, the samples were similar. Figure 2 shows sections of microphotographs with margins of approximately 150 μm from three typical examples 3.1. Subsection

**Figure 1.**
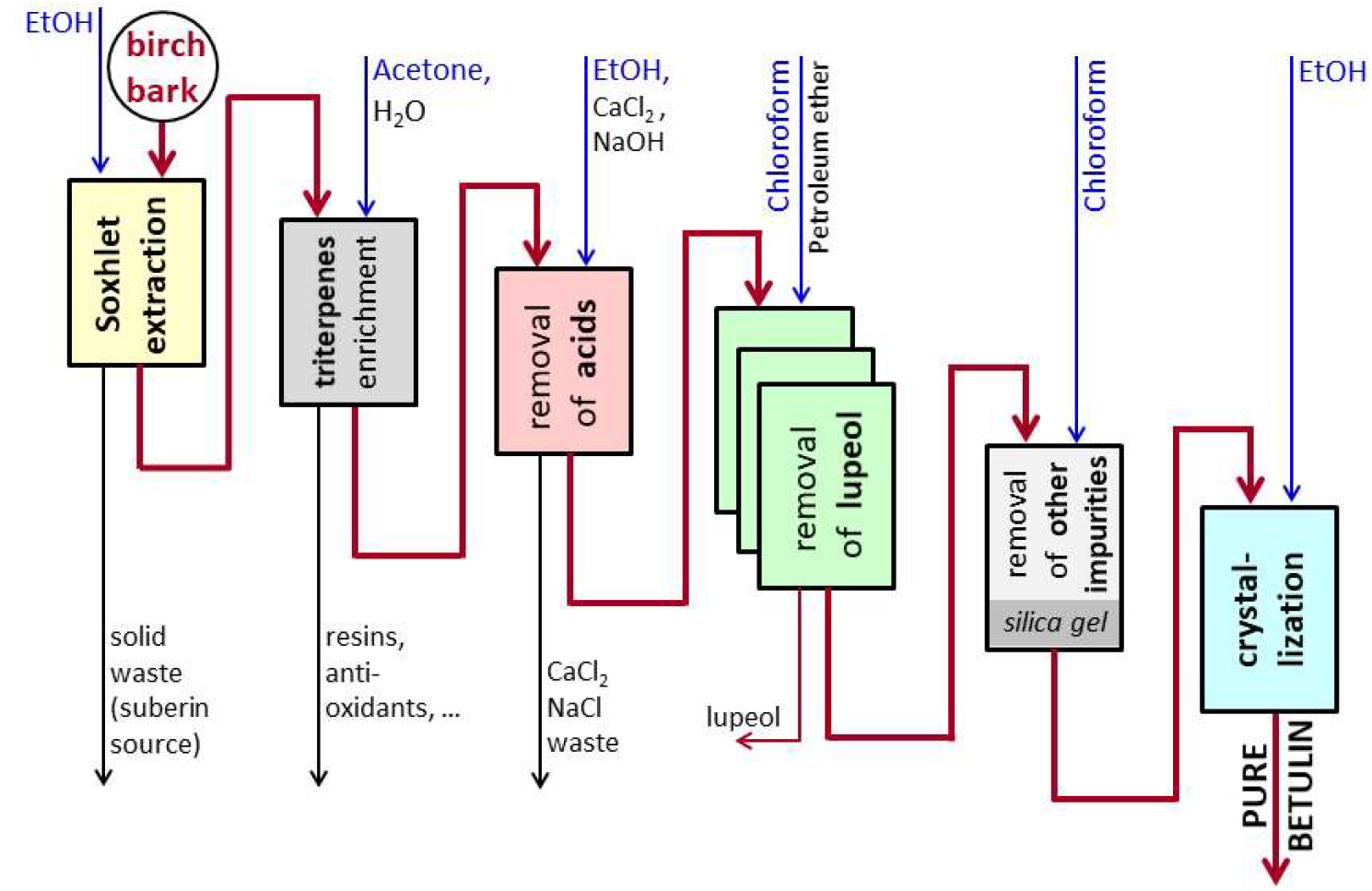
Scheme of pure betulin preparation (modified from Šiman et al. [5]).

**Figure 2.**
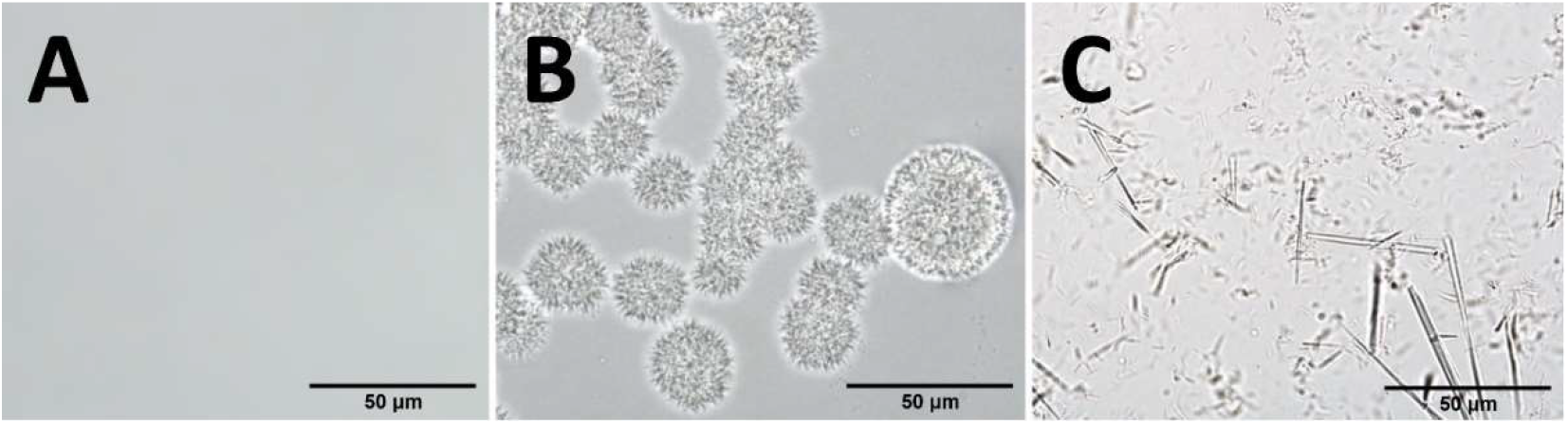
Photomicrographs of betulin in carriers. (**A**) Olive oil (10 mg/ml); (**B**) olive oil (16 mg/ml); (**C**) cocoa butter (4 mg/ml). The selected photos were taken within a maximum of 6 hours after the preparation of the solutions.

### 3.2. Administration of betulin per os

Betulin administration using lard in feed seemed problem-free. Standard feed enriched in lard tasted good to the rats. Unfortunately, we did not measure the consumption of enriched feed in comparison with standard feed.

Both methods of betulin administration, with lard in feed and via gavage as a solution in olive oil, were successful. This finding was confirmed by measuring betulin levels in the organs of the rats and the visible effect on the mass of the solid tumors in the mice together with Namitecan, as shown below.

The method used indicated an order of magnitude greater availability and 2-3 orders of magnitude greater efficiency (considering the amount of betulin administered) of the given form of betulin solution than the use of conventional suspensions for *per os* or intraperitoneal administration [14,15].

### 3.3. Administration of betulin per rectum

The administration of betulin *per rectum* using a suppository mass as a carrier did not provide the desired results. Although some detectable amounts of betulin appeared in the jejunum 5 hours after application, the level in whole blood remained below the detection limit of the method.

### 3.4. Determination of betulin concentrations in organs

The method for determining betulin in organs is simple to perform and sensitive enough to reliably determine therapeutically appropriate amounts of betulin (e.g., 5 μg/g according to the IC50 in the case of the triple-negative breast tumor cell line BT-549 - Šiman et al. [5] or, in the case of some other cell lines, Król et al. [1]). The authors estimate the sensitivity limit to be much less than 1 μg/g tissue when GC-MS is used. Heparinized blood appears to be the best among all organs tested.

Only two rats were used to determine the distribution of betulin administered *per os*. Therefore, the “statistics” represented by the “standard deviation” in the following figure are highly questionable. Nevertheless, notably, there were very few differences in all the organs studied between the two rats, with the exception of adipose tissue. Thus, it appears that the two animals were fed identically with respect to their weight and that the equilibrium distribution of ingested betulin in the organs was quite characteristic. The adipose tissue undoubtedly contained a considerable amount of betulin, but processing of the samples by the method described was unsuitable for this tissue. The high fat content led to the formation of large amounts of sodium soap, which precipitated as a shapeless mass containing an unknown amount of betulin not available for extraction to chloroform. For adipose tissue, therefore, the methodology would have to be modified.

The next figure shows substantial differences among various organs (up to sevenfold between kidney and blood sediment), especially compared with small individual “standard deviations”, i.e. the simple differences between the both obtained values.

### 3.5. Synergistic effects of Namitecan and Betulin

The results of this pilot study on the effect of betulin in cancer treatment are promising. Although betulin alone is not effective (P = 0.0361) in the *in vivo* treatment of Ehrlich’s adenocarcinoma (Figure 4), as reported in some tumor types even *in vitro* [1], it can act synergistically with other cytostatic drugs. The cytostatic drug Namitecan showed, as expected, high treatment efficiency (P < 0.001). However, the combination of Namitecan with betulin showed synergistic effects, and an even greater efficiency was achieved in the *in vivo* treatment of Ehrlich’s adenocarcinoma than in the control (P < 0.001) and Namitecan alone (P = 0.024) (Figure 4).

**Figure 3.**
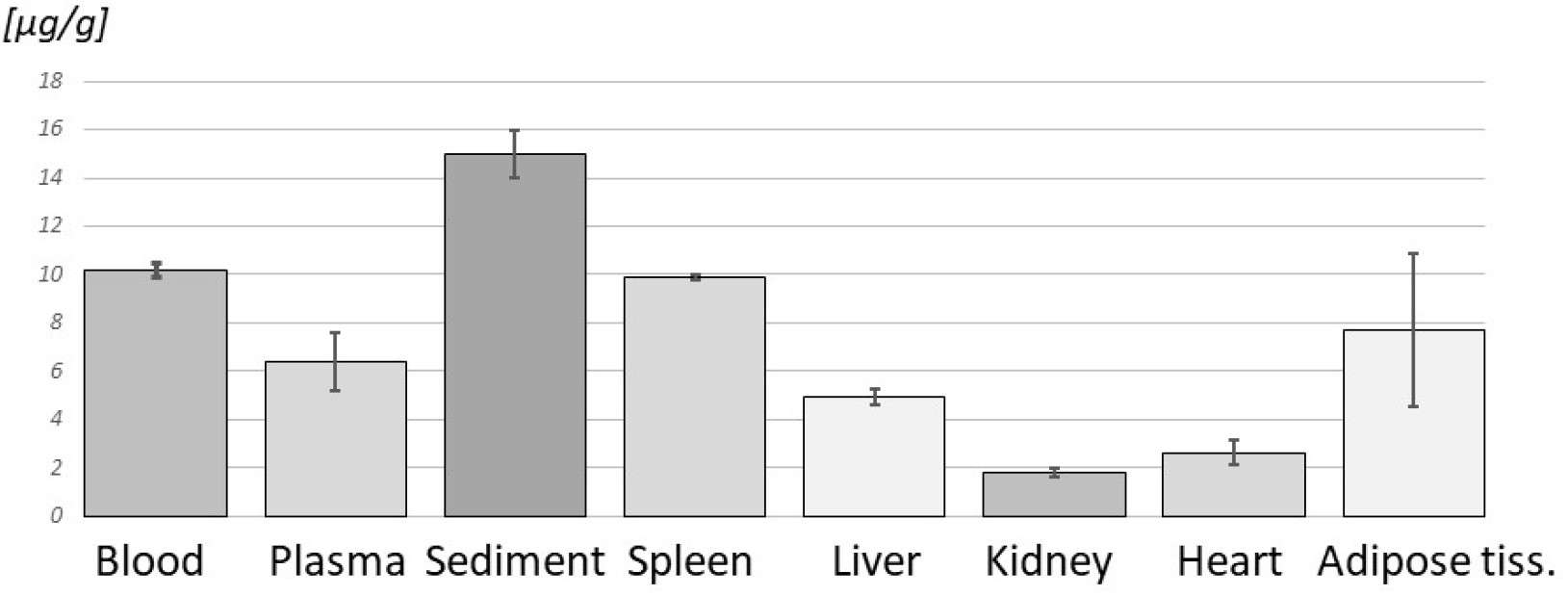
Betulin in rat tissues. Levels of betulin after one week of administration of lard with betulin (μg of betulin/g tissue; n=2).

**Figure 4.**
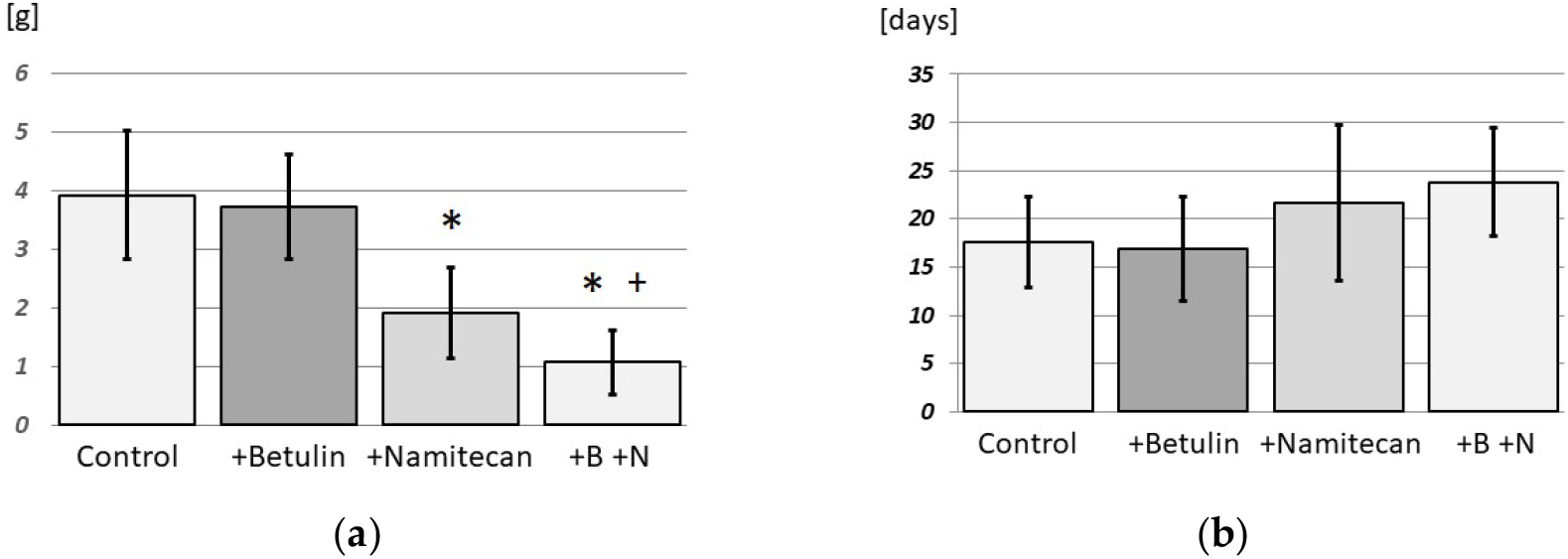
(**a**) Mass of solid Ehrlich’s tumor after treatment with betulin and/or Namitecan; (**b**) Namitecan and survival of treated mice. n=7 in control, betulin, and Namitecan; n=6 in N+B. The asterisk (*) denotes significantly lower Ehrlich’s tumor weight in comparison with the control group (+Namitecan; +B +N; P < 0.001). The plus sign (+) denotes significantly lower Ehrlich’s tumor weight than the Namitecan group (P = 0.024).

## 4. Discussion

### 4.1. Carriers for betulin

The acylglycerols used were relatively good carriers of molecular betulin. In particular, olive oil allows the uptake of therapeutic amounts of betulin by organisms. The lard can also be used as part of a mixed diet (but it is not well defined as a carrier), since the rats seemed to like an enriched diet. However, it would be extremely difficult to obtain true solutions by merely dispersing pure betulin in given carriers, since crystalline betulin dissolves very slowly even in hot ethanol, for example. In addition, betulin dispersed by direct dissolution does not have to be in molecular form, as even its nanoclusters can externally appear as a real solution owing to transparency. The same will be the case with the use of suspensions of any size of betulin particles, whether they are used orally, subcutaneously, by inhalation or even intraperitoneally. Compared with a purely molecular form, there can be a fundamental difference in the amount of betulin taken up by the organism. The chosen method of indirect dissolution, i.e., introducing the ethanol solution into the carriers and then evaporating the ethanol, proved to be extremely suitable and effective. Since betulin is in its molecular form in the ethanolic solution, there is no reason to assume that it will subsequently form nanoclusters in the carrier. Rather, and this was demonstrated microscopically, it simply crystallizes in microcrystals when its solubility in carriers is exceeded.

Evaporation of the ethanol from the carriers could be further improved by reducing the pressure while inert gas bubbles pass through (N_2_, Ar, e.g.). The carriers used were not pretreated in any way. However, it is conceivable that they could largely absorb betulin after thorough drying, for example. For such drying, inert gas bubbling through a liquid warm carrier up to 100 °C with reduced pressure, e.g., could be suitable for the pharmaceutical production of cachets with betulin-enriched olive oil.

The use of a simple ethanol solution of betulin placed directly on food, i.e., without the use of a proper carrier [22], is highly questionable. This approach is based on the authors’ experience with betulin solutions in various solvents. The ethanol evaporates, and betulin precipitates as microcrystals. Thus, the availability of betulin to experimental animals is extremely limited, and it would be difficult to detect betulin in experimental subjects and achieve reproducible results.

The proposed method of administration may lead to revision of some quantitative data from *in vivo* experiments, especially in the case of pharmacokinetics [4] and safety or lethal doses of triterpenoids [7].

### 4.2. Determination of betulin concentrations in organs

Most *in vivo* studies avoid determining the levels of betulin in the organism, as such determination places high demands on objectivity. The method described in, e.g., Jäger et al. [14], is controversial, and it is difficult to deduce the real betulin level from it, as the following lines of discussion show.

For the determination of betulin in tissues, it is necessary not only to dissolve the tissue but also to saponify fats and hydrolyze all polymers with hydrophobic components, e.g., proteins. Only then can it be assumed that all betulin is available for chloroform extraction. The solution to the aforementioned problems is to use a high concentration of NaOH, which hydrolyzes most of the polymers presented and converts the fats and fatty acids into the ionic form of soaps. Both betulin and lupeol are stable even at this high pH. Since extraction with chloroform is carried out at laboratory temperature and over a relatively short period of time, there is practically no risk of a noticeable reaction between chloroform and NaOH.

However, the method used for the hydrolysis of polymers and saponification of fats with high NaOH concentrations is unlikely to be suitable for accessing acidic triterpenoids such as betulinic acid, oleanolic acid or ursolic acid. Here, it is virtually certain that they are ionized at high pH and cannot be quantitatively extracted into chloroform.

Only two rats were used for the determination of betulin in the organs; therefore, no statistically proven conclusions could be drawn. However, the very small variations in the values obtained, with the exception of adipose tissue, where massive precipitation of soaps was probably the main cause, suggest at least sufficient distribution of betulin administered *per os* among the different organs after one week. This finding also suggests which organ tumors sensitive to betulin are most affected. Given that the very small deviations between the organs of the two rats can hardly be the work of a happy accident, the results also demonstrate the high reliability of the proposed method for determining betulin in organs.

Since there are no studies on the basis of which the appropriate time of administration could be determined, this was determined by estimation to establish an equilibrium concentration of betulin in the rats. Importantly, this study is the first pilot study to create an effective form of betulin administration, in which betulin remains safe in the molecular form of the true solution and is therefore absorbable by the organism. This study is also the first to provide a method for determining true betulin levels in organs. Only now will it be possible to carry out further necessary experiments, e.g., pharmacokinetics of betulin, histological control of basic vital organs, and monitoring of the overall condition of the animals after longer-term exposure to betulin, which will enable a qualified determination of the appropriate administration time.

The determination of betulin in complete heparinized blood could reflect the level of biologically available betulin in the body. On the one hand, there is a significant amount of betulin in the blood, and on the other hand, the blood can be taken easily during the entire period of betulin administration. For example, pharmacokinetic studies can be performed relatively easily. In clinical trials, this is also the only easy way to monitor betulin levels in the body during treatment. Blood lipoproteins are likely the main carriers of betulin in the body.

An attempt has also been made to determine betulin in clotted blood, but this has not been successful. Initially, it was quite difficult to dissolve blood clots. Such samples were then often strongly discolored in the chloroform layer and thus had to be removed from samples for measurement by GC-MS. Therefore, it is far better to use blood that has not clotted (heparinized, e.g.).

Fifteen males of NMRI mice were used for the optimization of individual methodologies, e.g., administration processes and experiments with clotted blood, without relevant experimental data.

However, important information emerged from these experiments, such as the inappropriateness of using clotted blood to determine the content of betulin in the body.

The determination of betulin in the organs of the mice was not planned in this study because of the use of a very low concentration of betulin in the applied olive oil and the resulting assumption of the probable disappearance of betulin in the organs of the mice at a time shorter than their expected survival time.

### 4.3. Synergistic effects with Namitecan

The results of this pilot study on the synergistic effect of betulin with other cytostatic drugs in tumors insensitive to betulin alone are promising. The synergistic effect of using Namitecan with betulin in the *in vivo* treatment of Ehrlich adenocarcinoma was demonstrated, as the group treated with Namitecan with betulin presented significantly lower Ehrlich tumor weights than did the group treated with Namitecan alone. The difference in survival times was not statistically significant in this pilot experiment; however, the results indicated a positive trend (Figure 4). It is very likely that very good results could be obtained after appropriate modification of the experimental protocol.

Notably, only a very small amount of betulin was used, and its concentration in the administered olive oil could be increased up to 20-fold. Betulin could also be administered for a longer period than it was in the experiment. All this could have significantly changed the survival of the experimental animals. In the case of strong statistical evidence of the synergistic effect in subsequent full-scale experiments, the use of betulin may reduce the doses of cytostatic drugs, reducing their side effects.

Moreover, betulin administered together with a more aggressive cytostatic drug may also have a protective effect on organs, as suggested, for example, in Lee et al. [23], in which triterpenes showed a renoprotective effect when cis-platinum was used.

### 4.4. Stability of pure betulin

The crystalline form and purity of highly pure betulin did not change, as detected microscopically (Figure 5) and by GC-MS, even after five years of storage in the dark in a well-closed container and at temperatures ranging from approximately 15–40 °C. The recommended storage times, especially the expiry times, of commercial products are therefore often excessively strict. However, this may also be related to purity, as the betulin used was extremely pure (99.7%) according to GC-MS analysis.

**Figure 5.**
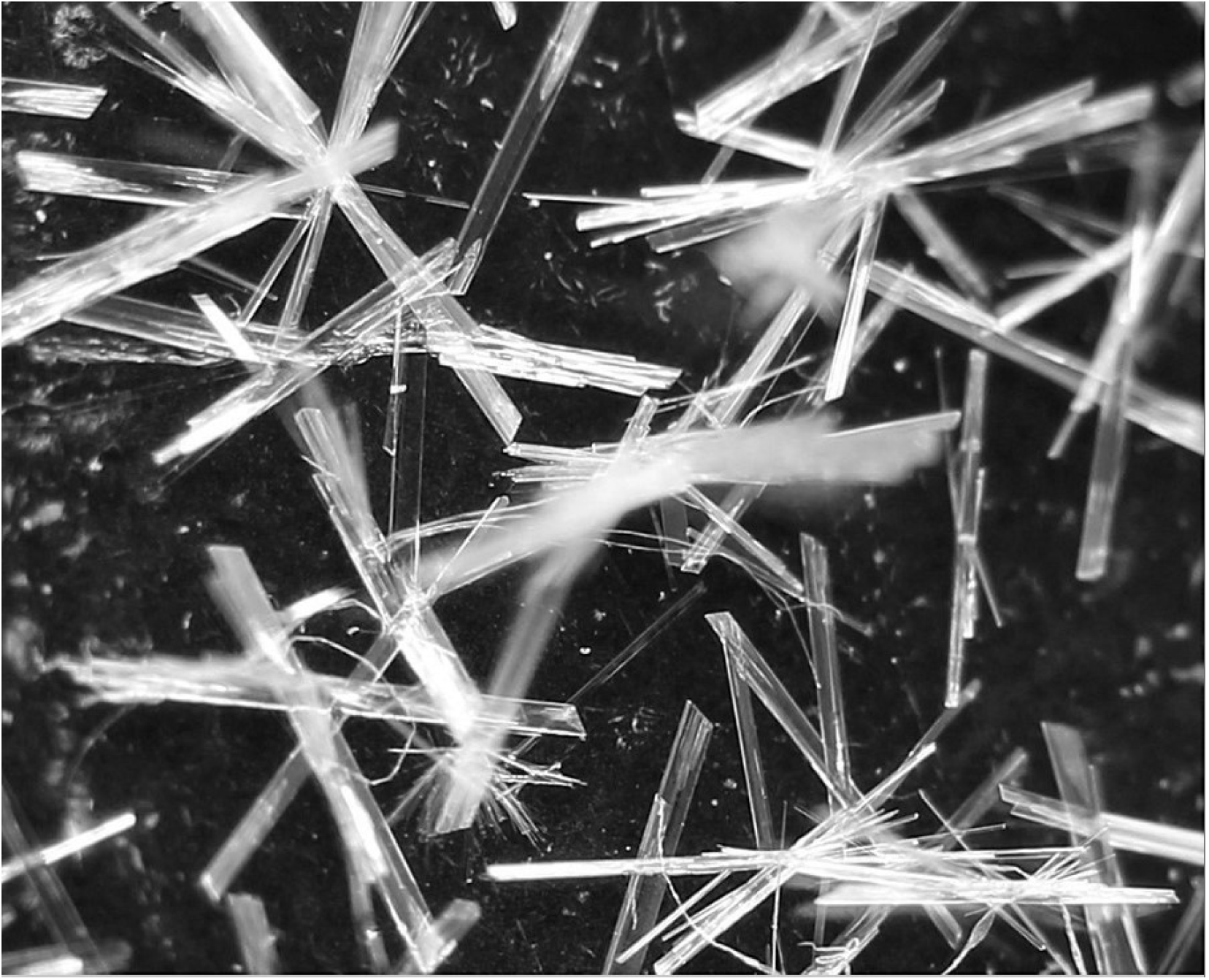
Pure betulin five years after preparation.

### 4.5. Microbial stability

The microbial stability of betulin is sufficient, because it is sterile after final crystallization from ethanol in its preparation by the method we used and as a solid crystalline substance it can be stored sterile. However, this is not necessary, because during the preparation of the dosage form betulin again enters in the form of a hot ethanol solution. The resulting solutions in acylglycerols can be easily subsequently sterilized by heat. In addition, as mentioned in the literature, betulin itself exhibits certain antimicrobial, antiviral and antifungal effects.

### 4.6. Strengths and limitations

The method of preparing betulin for *per os* application enables the generation of a true/molecular solution of betulin in the selected carrier with a completely reproducible composition and reproducible bioavailability for the organism under investigation. This will now allow further *in vivo* experiments to be evaluated truly quantitatively, not just qualitatively in nature. The second method, the analysis of samples, enables continuous and reproducible determination of the actual levels of betulin in the organism under investigation.

All the preliminary observations mentioned above need to be confirmed by further research and on a larger scale. However, the promising and statistically significant results from this pilot small-scale experiments provide hope for further full-scale studies on the anticancer effects of betulin and for preclinical and clinical studies.

Unfortunately, the authors can no longer continue their work. However, they believe that their pilot work can be the basis for research by other scientific groups with the ultimate goal of successful clinical trials using the natural substance betulin, an inexpensive drug with an almost infinite source in birch bark.

## 5. Conclusions

The introduction of a hot ethanol solution of betulin to warm acylglycerols seems to be the best method to obtain a true solution of betulin in carriers such as olive oil, suppository mass or lard *per os* administration of this triterpenoid in *in vivo* experiments. The concentration of betulin in such carriers may be sufficient for reaching therapeutic levels in experimental animals or, in the future, in the bodies of potential patients. This method of administration is gentle to animals or patients and is inexpensive, relatively quick and effective with the delivery of betulin to the body. The suggested method of administration is supplemented by a relatively simple analytical method for the determination of betulin in organs, especially in the blood. The samples were dissolved in NaOH solution, and then betulin was extracted with chloroform. The content of betulin in the evaporated residue of the chloroform solution was then analyzed chromatographically.

The present method of administering betulin proved to be suitable for studying its synergistic action with the cytostatic drug Namitecan. Although this study represents the first pilot experiment and the amount of available betulin was very small, the results proved to be promising, and the synergistic effect was statistically proven.

## Author Contributions

Writing – original draft: P.Š. and A.B.; Writing – review & editing: P.Š., A.B., A.T., H.K., T.H., O.K. and M.N.; Conceptualization: P.Š. and M.N.; Data curation: P.Š., A.B., and A.T.; Formal analysis: A.B., P.Š., and A.T.; Investigation: P.Š., A.B., A.T., H.K., T.H., O.K. and M.N.; Methodology: P.Š.; Resources: P.Š., O.K. and M.N.; Supervision: P.Š.; Project administration: P.Š.; Validation: P.Š. and A.B.; Visualization: P.Š. and A.B.

## Institutional Review Board Statement

All animals were handled according to the guidelines of the Institutional Animal Use and Care Committee of Charles University, Prague, Czech Republic, and in accordance with the recommendations of the Animal Care Committee of the Institute of Microbiology of the Czech Academy of Sciences (approval ID: 31/2020). The Institute of Microbiology of the Czech Academy of Sciences was granted permission to use experimental animals in relation to this facility, file number 16OZ11877/2019-18134, case number 37330/2019-MZE-18134, valid from 18 August 2019 to 15 August 2024. Furthermore, RNDr. Hana Kozáková was granted a certificate of professional competence for designing experiments and experimental projects under Section 15e (1) of Act No 24611992 Coll., on the Protection of Animals against Cruelty No. CZ 01521 valid from 27 April 2021 to 27 April 2028.

## Informed Consent Statement

Not applicable.

## Data Availability Statement

The data that support the findings of this study are available from the corresponding author upon reasonable request.

## Acknowledgments

The article is dedicated to the memory of dr. Mohamed Niang.

## Abbreviations

The following abbreviations are used in this manuscript:

GC-MS: gas chromatography with mass spectrometry
UHPLC-MS: ultra-high performance liquid chromatography with mass spectroscopy IC50 half maximal inhibitory concentration

